# Similar ratios of introns to intergenic sequence across animal genomes

**DOI:** 10.1101/068627

**Authors:** Warren R. Francis, Gert Wörheide

## Abstract

One central goal of genome biology is to understand how the usage of the genome differs between organisms. Our knowledge of genome composition, needed for downstream inferences, is critically dependent on gene annotations, yet problems associated with gene annotation and assembly errors are usually ignored in comparative genomics. Here we analyze the genomes of 68 species across 12 animal phyla and some single-cell eukaryotes for general trends in genome composition and transcription, taking into account problems of gene annotation. We show that, regardless of genome size, the ratio of introns to intergenic sequence is comparable across essentially all animals, with nearly all deviations dominated by increased intergenic sequence. Genomes of model organisms have ratios much closer to 1:1, suggesting that the majority of published genomes of non-model organisms are underannotated and consequently omit substantial numbers of genes, with likely negative impact on evolutionary interpretations. Finally, our results also indicate that most animals transcribe half or more of their genomes arguing against differences in genome usage between animal groups, and also suggesting that the transcribed portion is more dependent on genome size than previously thought.

## Introduction

Understanding why genomes vary greatly in size and how organisms make different use their genomes have been central questions in biology for decades [1]. For many bacteria, the majority of the genome is composed of relatively short genes, averaging around 1000bp, and coding for proteins. Indeed, the largest bacterial genome (a myxobacterium) that has been sequenced is only 14 megabases, containing an estimated 11,500 genes [2]. However, for eukaryotic organisms, genomes can be over ten-thousand-fold larger than bacterial genomes due to an increase in the number of genes (tens of thousands compared to a few thousand in most bacteria), expansion of the genes themselves due to the addition of introns, and expansion of the sequence between genes.

As the number of genome projects has grown, massive amounts of data have become available to study how organisms organize and use their genomes. Genome projects vary substantially in quality of assembly and annotation [3, 4]. Unfortunately, the predicted genes are often taken for granted as being correct when these are only hypotheses of gene structure [5]. For example, one study found that almost half of the genes in the *Rhesus* monkey genome had a predictable annotation error when compared to the closest human homolog [6]. This has profound implications for all downstream analyses, such as studying evolution of orthologous proteins [7] and phylogeny based on protein matrices or gene content [8, 9]. When considered across all genes, systematic errors in genome assembly or annotation would severely skew bulk parameters of a genome.

While issues of assembly are often thought to be technical problems that are resolved before continuing, all subsequent analyses are dependent upon accurate genome assembly and annotation. The absence of a protein family in a particular organism is only meaningful if it is certain that it is absent from the genome and not merely the annotation, therefore it is of utmost importance that all genes are properly represented. Yet for most genome projects of non-model organisms, there are limited methods to determine if the assembly and annotation are sufficient for downstream comparative analyses. Internal metrics can be used, such as the fraction of raw genomic reads or ESTs that map back to the assembly, though this does not tell us if a gene is believable in the context of other animals. Alternatively, counts of “universal” single-copy orthologs have been proposed as a metric of genome completeness [10, 11], though these genes only represent a small subset of all genes (few hundred out of tens of thousands in most animals).

Identification of universal trends in genome organization and transcription may enable better quantitative metrics of genome completeness. Mechanistic models relating to evolution of gene content or coding fractions tended to focus on bacteria or archaea because of the relative ease of annotation. In regards to eukaroytes, some patterns in genome size have been discussed [12–14]. Additionally, a handful of studies have analyzed genome size in connection to other parameters such as indels [15], transposon content [16–19], average intron length [20, 21] or total intron length [18]. Despite these advances, none of these studies have estimated the amount of the genome that is genic (exonic plus intronic, including non-coding) based on independent examination of single genomes and without averaging over a whole kingdom. Additionally, none of them have described a way to account for technical problems in assembly and annotation.

Here we examine basic trends of genome size and the relationship to annotation quality across animals and some single-celled eukaryotes. We show that assembly and annotation errors are widespread and predictable and that many genomes are likely to be missing many genes. We further show that re-annotation of select species with publicly available tools and transcriptome data improves the annotation. Future users may benefit if databases incorporate more recent data from transcriptome sequencing, and update annotation versions more frequently. Comparison of genomic composition across many animal groups indicated a ratio of introns:intergenic approaching 1:1, suggesting this as a potential parameter to identify genome completeness across metazoans, and potentially other eukaryotes. Finally, this implies that animals transcribe at least half of their genomes whereby small, exon-rich genomes transcribe most of the genome and large genomes transcribe approximately half of the genome.

## Methods

### Genomic data sources

#### Data sources and parameters are available in Supplemental Table 1

Genomic scaffolds and annotations for *Ciona intestinalis* [22], *Branchiostoma floridae* [23], *Trichoplax adherens* [24], *Capitella teleta* [25], *Lottia gigantea* [25], *Helobdella robusta* [25], *Saccoglossus kowalevskii* [26], *Monosiga brevicollis* [27], *Emiliania huxleyi* [28], and *Volvox carteri* [29] were downloaded from the JGI genome portal.

Genome assemblies and annotations for *Sphaeroforma arctica*, *Capsaspora owczarzaki* [30] and *Salpingoeca rosetta* [31] were downloaded from the Broad Institute.

GFF annotations v2.1 [32] for *Amphimedon queenslandica* were downloaded from the Amphimedon Genome website (http://amphimedon.qcloud.qcif.edu.au/downloads.html), and v1 annotations [33] and assemblies were downloaded from Ensembl.

For *Nematostella vectensis*, Nemve1 assembly and annotations [34] were downloaded from JGI, and the transcriptome for comparative reannotation was downloaded from http://www.cnidariangenomes.org/ [35].

Genome assembly, transcriptome assemblies from Cufflinks and Trinity, and GFF annotations for *Mnemiopsis leidyi* [8] were downloaded from the Mnemiopsis Genome Portal (http://research.nhgri.nih.gov/mnemiopsis/). Assembly and annotations for *Sycon ciliatum* [36] were downloaded from COMPAGEN. Assembly and annotation for *Botryllus schlosseri* [37] were downloaded from the Botryllus Schlloseri genome project (http://botryllus.stanford.edu/botryllusgenome/). Assembly and annotation for *Exaiptasia pallida* (formerly *Aiptasia sp.*) [38] were downloaded from http://reefgenomics.org Assembly and annotation for *Oikopleura dioica* [39] were downloaded from Genoscope (http://www.genoscope.cns.fr/externe/GenomeBrowser/Oikopleura/). Assembly and annotation for *Tetrahymena thermophila* were downloaded from the Tetrahymena Genome Database (ciliate.org). Assembly and annotation for *Symbiodinium kawagutii* [40] were downloaded from the Dinoflagellate Resources page (web.malab.cn/symka new/index.jsp).

Assemblies and annotations for *Symbiodinium minutum* [41], *Pinctada fucata* [42], *Acropora digitifera* [43], *Lingula anatina* [44], *Ptychodera flava* [26], and *Octopus bimaculoides* [45] were downloaded from the OIST Marine Genomics Browser (http://marinegenomics.oist.jp/gallery/).

Builds of *Homo sapiens*, *Pan troglodytes*, *Mus musculus*, *Canis lupus* [46], *Monodelphis domestica* [47], *Ornithorhynchus anatinus* [48], *Xenopus tropicalis* [49], *Struthio camelus* [50], *Gallus gallus*, *Taeniopygia guttata* [51], *Aptenodytes forsteri* [50], *Anas platyrhynchos* [52], *Melopsittacus undulatus* [53], *Alligator mississippiensis* [54], *Anolis carolinensis* [55], *Chrysemys picta bellii* [56], *Chelonia mydas* [57], *Pelodiscus sinensis* [57], *Python bivittatus* [58], *Salmo salar*, *Danio rerio* [59], *Latimeria chalumnae* [60], *Petromyzon marinus* [61], *Callorhinchus milii* [62], *Crassostrea gigas* [63], *Dendroctonus ponderosae* [64], *Tribolium castaneum* [65], *Bombyx mori* [66], *Limulus polyphemus* [67] were downloaded from the NCBI Genome server.

Genome assemblies and annotations of *Caenorhabditis elegans* [68], *Drosophila melanogaster*, *Strongylocentrotus purpuratus* [69], *Daphnia pulex* [70], *Apis mellifera* [71], *Ixodes scapularis* [72], *Strigamia maritima* [73] were downloaded from Ensembl.

### Calculation of exonic and genic sequence

For all analyses, we used the total number of bases in the downloaded assembly as the total genome size, bearing in mind that this may result in a systematic underestimation of total genome size as repeated regions may be omitted from assemblies. For example, the horseshoe crab *L. polyphemus* has a scaffold assembly of 1.8Gb while the reported genome size is 2.7Gb [67], a difference of almost a gigabase.

If GFF format files were available for download with a genome project, or on databases (Ensembl or NCBI), those were used preferentially. The analysis procedure is described in Fig 1. Total base pairs of exon, intron, intergenic, and gaps were counted from each GFF file and genomic contigs (or scaffolds) with a custom Python script (gtfstats.py, available at bitbucket.org/wrf/sequences). For calculations of exonic or genic bases, the script converts all gene and exon annotations to intervals and ignores the strand. Here, gene (or genic) is defined as transcribed bases that are either exon or intron, regardless of coding potential. All overlapping exon intervals are merged, meaning that alternative splice sites, or exons on the opposite strand, are treated as a single interval for bulk calculations. The same is done for genes or transcripts, whichever is available. Introns are calculated as the difference of the genic set and the exonic set, as introns are typically not defined as separate features in normal GFF files. This means that any sequence that is an exon on one strand and an intron on the other is treated for these calculations as an exon, meaning those base or their reverse complement (hence base pairs) are transcribed and retained following splicing in some case (Fig 1D and E). Intergenic sequence is defined as the difference between total sequence base pairs and genic base pairs, and gaps are defined as any repeats of ‘N’s longer than one base.

**Figure 1:**
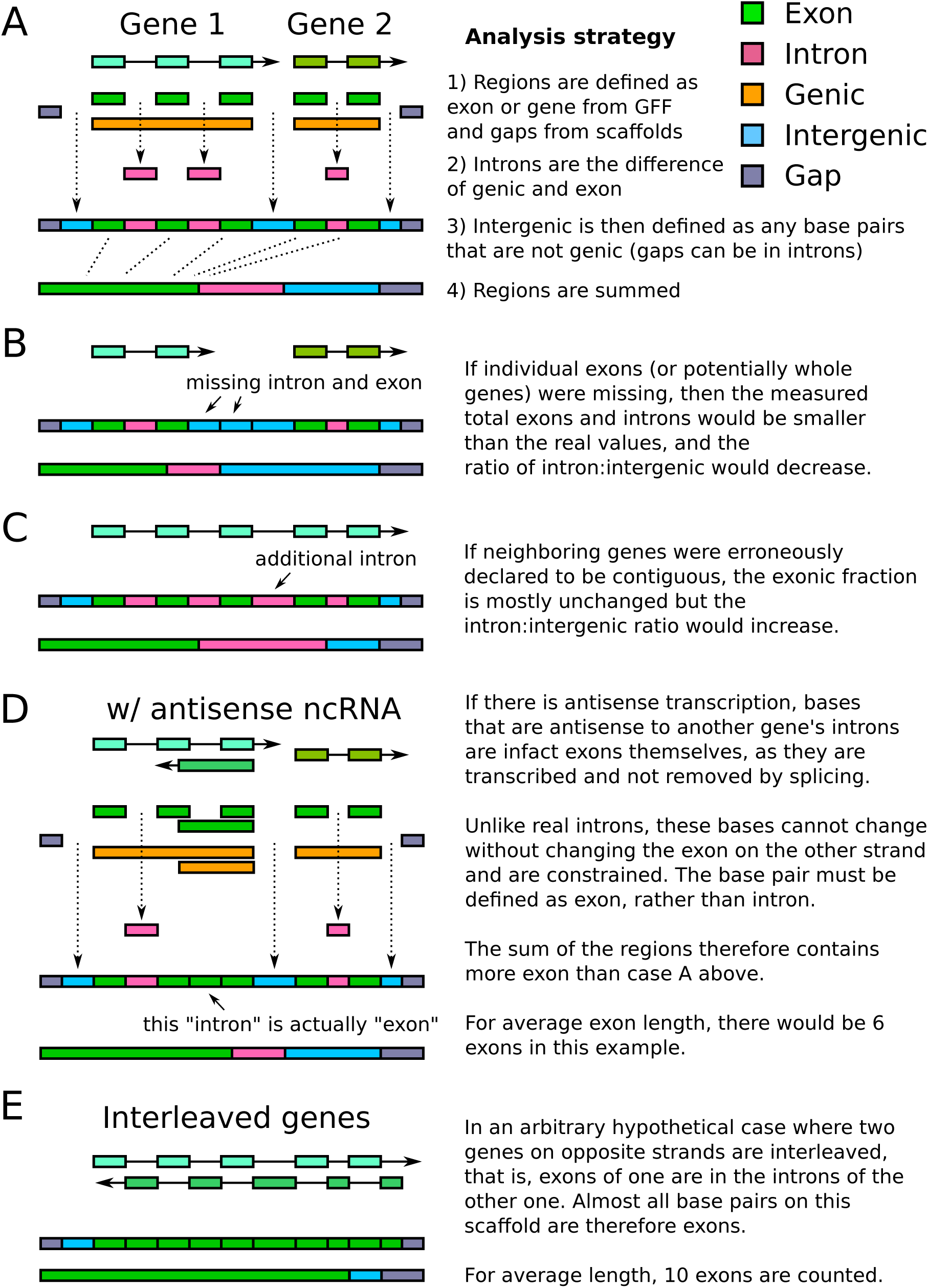
Schematic of analysis, misannotations and the effects on coding fraction. (A) In a normal case, two hypothetical genes on the same strand are identified. The exons and introns are defined, and the total lengths of those features are summed and display7ed in the bars below. Because real genome assemblies can often contain gaps, sample gaps are also shown at the edges of the segment. (B) Case of missing exon or gene annotations, where the intron:intergenic decreases. (C) Case of falsely fused genes, where the intron:intergenic ratio would increase. (D) Case of antisense transcription, where base pairs that are intron on the sense strand and exon on the antisense strand are necessarily defined as exon. (E) Any arbitrary, interleaved genes, or any exons inside of introns, must as well be counted as exon.

If exons are not specified, then coding sequences (CDS) are used instead if they are available, such as for AUGUSTUS predictions. Additional non-coding features such as “microRNA”, “tRNA”, “ncRNA” are included for gene and exon calculations if they were in the standard GFF3 format. Some genomes made use of mapped RNAseq data, which implicitly included all non-coding RNAs as well. Some annotations had to determine the gene ID from the exons. For example, most of the GTF files from the earlier JGI genomes had only exons annotated, without individual features for genes or mRNAs, so the gene was then defined as all of the exons with the same feature ID even though a specific gene feature was undefined.

Exons defined as part of a “pseudogene”, or genes defined as pseudogenes, were also excluded from all counts. We justify this because pseudogenes are subject to problems of definitions and population sampling bias. Pseudogenes are defined as having the appearance or structure of normal protein coding genes, independent of transcriptional potential, but that would be unable to produce a functional protein, perhaps through nonsense mutations. Therefore, a pseudogene that is transcribed and cannot code for a protein should be annotated as a “transcribed pseudogene”, though potentially could be a non-coding RNA. Pseudogene features are not annotated for all species, making it difficult to compare broadly. Additionally, for most non-model species, the genomes are generally based upon a single individual rather than a reference for a population based on a large number of individuals. Therefore, if that single individual were homozygous for a nonsense mutation but other individuals in the population were not, that gene should not be a pseudogene.

All downstream correlation calculations and graphs were done in R. Regression was calculated using the “lm()” function, for linear (y*∼* x), exponential (log(y)*∼* x), or hyperbolic (y*∼*1/x) models, and the “predict()” function was used to model curves. The raw data table and the R source code used to generate figures is available at bitbucket.org/wrf/genome-reannotations.

### Calculation of average exon and intron length

The same script (gtfstats.py, available at bitbucket.org/wrf/sequences) also calculated the average exon and intron length, though these were analyzed separately. All non-redundant exons for all splice variants were taken into account for determination of averages. Unlike the total base pair calculations, genes are separated by strand. Identical exons of splice variants were treated as one exon and counted once, however, alternative boundaries were treated as a separate exons. Retained introns are treated as exons, not introns. Exon lengths were counted per non-redundant exon for each gene, summed across all genes and divided by the number of non-redundant exons across all genes. The sum of exon lengths for the average length calculation does include redundant bases from antisense transcripts or splice variants, meaning bases from antisense transcripts and alternative-boundary splice variants can be double-counted. Introns were calculated as the space between exons, calculated by gene.

### Reannotation of select species

Due to unexpectedly high or low gene content, six genomes were selected for reannotation.

The original Triad1 scaffolds of *T. adherens* [24] were reannotated with AUGUSTUS v3.0.3 [74] with the following options: -strand=both –genemodel=atleastone –sample=100 –keep viterbi=true –alternativesfrom-sampling=true –minexonintronprob=0.2 –minmeanexonintronprob=0.5 –maxtracks=2. Species training was generated using the Triad1 ESTs with the webAugustus Training server [75].

The original Monbr1 scaffolds of *M. brevicollis* [27] were reannotated with AUGUSTUS as for *T. adherens*, using the same parameters except trained using the Monbr1 ESTs with the webAugustus Training server [75].

For the hydrozoan *H. magnipapillata*, the original assembly was downloaded from JGI [76] and a new scaffold assembly was downloaded from the FTP of Rob Steele at UC Irvine (at https://webfiles.uci.edu/resteele/public). For both cases, the scaffolds were reannotated using TopHat22 v2.0.13 [77] and StringTie v1.0.4 [78] with default options by mapping the reads from two paired-end RNAseq libraries, NCBI Short Read Archive accessions SRR922615 and SRR1024340, derived from whole adult animals.

For the lancelet *B. floridae*, the Brafl1 scaffolds [23] were reannotated using TopHat22 v2.0.13 [77] and StringTie v1.0.4 [78] with default options by mapping the reads from the paired-end RNAseq library, NCBI SRA accession SRR923751, from the adult body.

For the lamprey *P. marinus*, we were unable to find any annotation as GFF or GTF, so we generated one using TopHat2 v2.0.13 [77] and StringTie v1.0.4 [78] based on the Pmarinus-v7 scaffolds from NCBI and the 16 single-end Illumina libraries from NCBI BioProject PRJNA50489.

For the octopus *O. bimaculoides*, scaffolds were downloaded from the OIST Marine Genomics platform [45], and were reannotated using TopHat2 v2.0.13 [77] and StringTie v1.0.4 [78] with default options by mapping 19 paired-end RNAseq libraries from NCBI BioProject PRJNA285380.

All reannotations are available for download as GTF or GFF files (see https://bitbucket.org/wrf/genomereannotations/downloads).

## Results

### Overview and organization of data

A total of 68 genomes were analyzed, with 59 selected across all major metazoan groups and nine genomes of single-celled eukaryotes. For each group, only select species were taken to avoid having a single group dominate the analysis. For example, over 100 mammalian genomes are available though only six were used including three model organisms (human, mouse, dog), opossum and platypus (for the non-eutherian clades, marsupial and monotreme, respectively) and the chimp, to compare directly to the human annotation. In general, parasites were excluded because they often have unusual biology, such as the single-celled eukaryote *T. brucei*, which is known for its unusual RNA processing [79, 80].

Generally, we refer to small and large genomes as those below and above 500Mb, respectively. The smallest animal genome used in this study is that of the larvacean *Oikopleura dioica* (70Mb), while the largest is that of the opossum *Monodelphis domestica* (3598Mb). This range incorporates an existing selection bias, as some of the public genome sequencing projects selected the animal of their clade based on their known small genomes. Two examples of this are the shark *C. milii* and the pufferfish *T. rubripes*. Yet it must be considered that in terms of genomes, they may not be representative of their clades; many other shark genomes are estimated to be over 10Gb (haploid genome size) [81], such that a shark genome of only 1Gb may not be “normal” for sharks.

Additionally, not all of the species in the sample were sequenced or annotated with the same method, making direct comparison more challenging. For instance, some of the earlier genomes (such as *Branchiostoma floridae* and *Trichoplax adherens*) were annotated only with Sanger ESTs (order of tens of Mb), which were used to train gene prediction algorithms. Because not all genes have features easily captured by the EST training, several different results are expected: some genes are split because internal exons are not properly found or may have misassemblies in the draft genomes; adjacent genes on the same strand are fused; or genes are omitted entirely.

### Connection between annotation and understanding of genomes

Genome projects of non-model species usually report protein coding regions of a genome. Broadly, there are two methods of doing this, comparison to other proteins from other genomes and by aligning mRNA from ESTs or RNAseq [3]. In practice, improvements in methods have made it relatively easy to directly predict proteins from the genome sequence. However, untranslated regions (UTRs) are difficult to predict and often require evidence from ESTs or transcriptome sequencing for accurate predictions, and this has implications for our measurements of total exons in each genome. This means that even in a “perfect” genome where all coding genes are correctly predicted by an annotation program (perhaps based on similarity to a related species) that the precise positions and amount of UTR may still be unknown, resulting in an underestimation of the amount of exonic sequence (Fig 1A and B). Because of this, the reliance on coding genes is likely to underestimate the usable fraction of the genome.

To illustrate this, one may consider a hypothetical eukaryotic genome of 60Mb with 10,000 genes and equal fractions of exons, introns, and intergenic sequence, at 20Mb each. For simplicity, all exons are the same size (in this example, 200bp), so an average gene (with ten-exons) may contain one exon for the 5’UTR, and one for the 3’UTR, and the remaining eight exons are coding. Based on the above annotation scheme, 20% of the exonic fraction (those containing the 5’ and 3’-UTRs) is missing in the final annotation. Two introns per gene are also missing (the first and last introns), about 18% of the intronic fraction. This would yield a final annotation where exons are predicted as 16Mb (26.6% of the genome) and introns as 15.5Mb (25.9% of the genome). This would also indicate that 52.6% of the genome is genes, a substantial underestimation from the actual value of 66.6%.

However, other systematic errors can result in an overestimation of the genic fraction. If we consider multiple genes on the same strand, in a head-to-tail arrangement, and recall that UTRs are often not predicted, then an exon containing the stop codon with a 3’-UTR may be omitted and the predicted gene may continue into the next gene (Fig 1C). If it is assumed that the majority of coding exons are correctly predicted, then if such predictions were made systematically one may expect that the measured amount of exons does not deviate much from the true exonic fraction. However, because introns are defined as the removed sequence between exons of the same gene, then the sequence between the two genes that should have been defined as intergenic will instead be defined as intronic, thus raising the intron:intergenic ratio above 1.

The above problems assume that the genomic assembly is nonetheless correct, yet the annotation is directly affected by assembly problems as well. Of the two main sources of problems, repeats [82] and heterozygosity [26, 42, 63, 83], repeats often result in breaks in the assembly that could split genes (Fig 2A). Genes that are split at contig boundaries are likely to have exons missing (or on other scaffolds) and thus the sequence that should be defined as introns would be instead defined as intergenic (Fig 2B).

**Figure 2:**
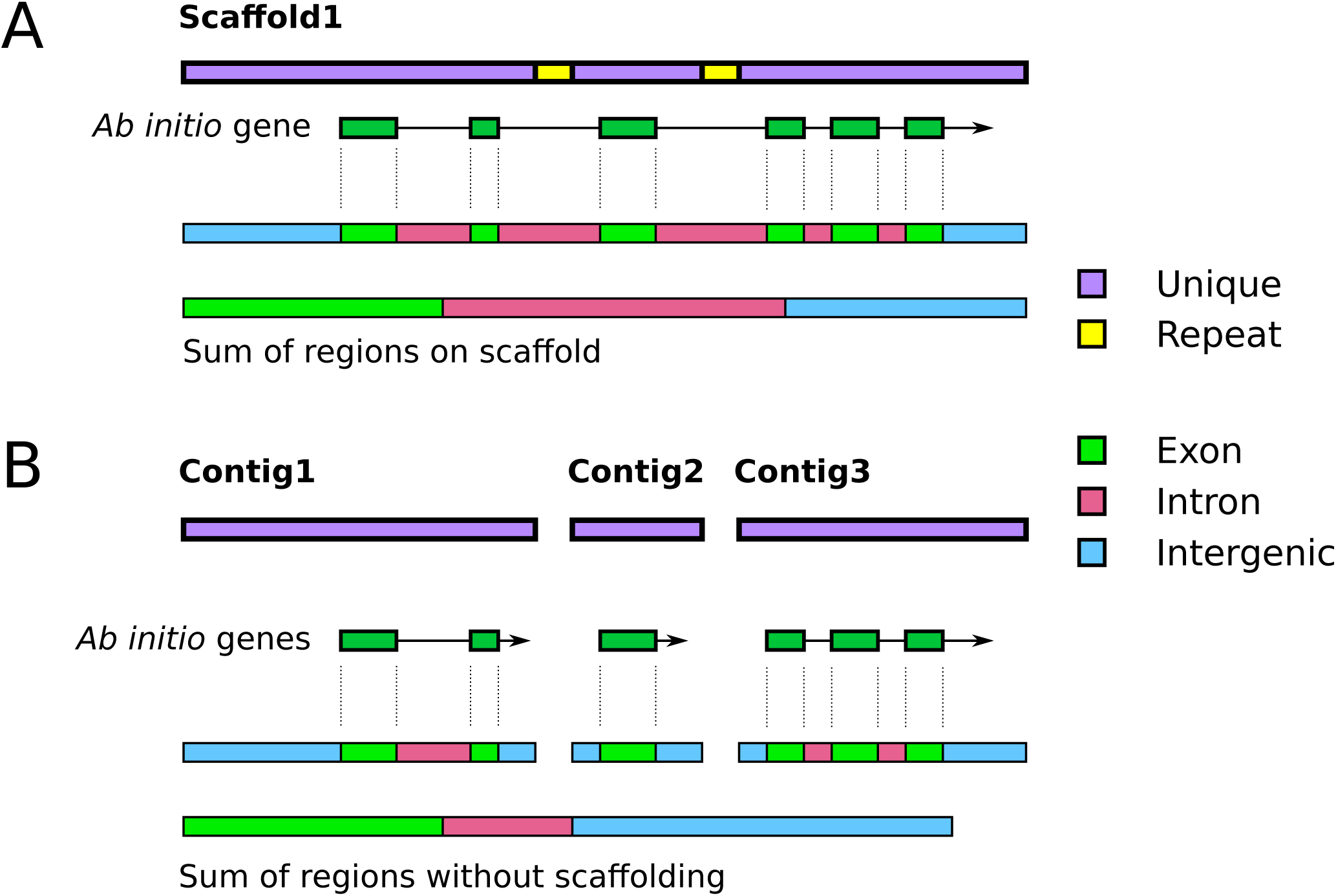
Schematic of the effects of scaffolding and repeats on genic fraction analyses. (A) For a hypothetical scaffold in a genome assembly, two identical repeats are found within introns. The gene is correctly predicted to span the two repeats and the regions are define below as in Fig 1. (B) For the case without scaffolding, or where the assembler breaks the assembly at repeats (or other high coverage regions), three contigs are generated. Note that the numbers are arbitrary, and in a real assembly they are unlikely to be in order. When annotated, all of the exons are correctly found, but the connections between them are missing for the single exon on Contig 2, resulting in a loss of intronic sequence. The final measured amount of exons is comparable, but the intron:intergenic ratio would decrease.

For normal diploid genomes (wild strains, not inbred lab strains), heterozygosity is not uniform across the genome. Some regions are identical between the two haplotypes (hence are homozygous alleles or loci), while others may vary by SNPs, short indels, or copy numbers of repeats, exons, or even genes. For sequences that are identical between both haplotypes, the contigs are generally kept as is, while a more complex decision must be made for the heterozygous loci. During normal genome assembly, the assembler evaluates the coverage at each “bubble” (where the de Brujin graph has two paths out of a node, and both paths merge again at the next node) and ultimately has to retain one of the paths at the exclusion of the other (Fig 3A) (also see schematics in [83] and [84]). This merging is the essential process that creates the reference genome, even though that reference is an arbitrary merge of the two haplotypes. Therefore, it must be kept in mind that predicted genes or proteins in reference genomes may not be identical to either haplotype.

**Figure 3:**
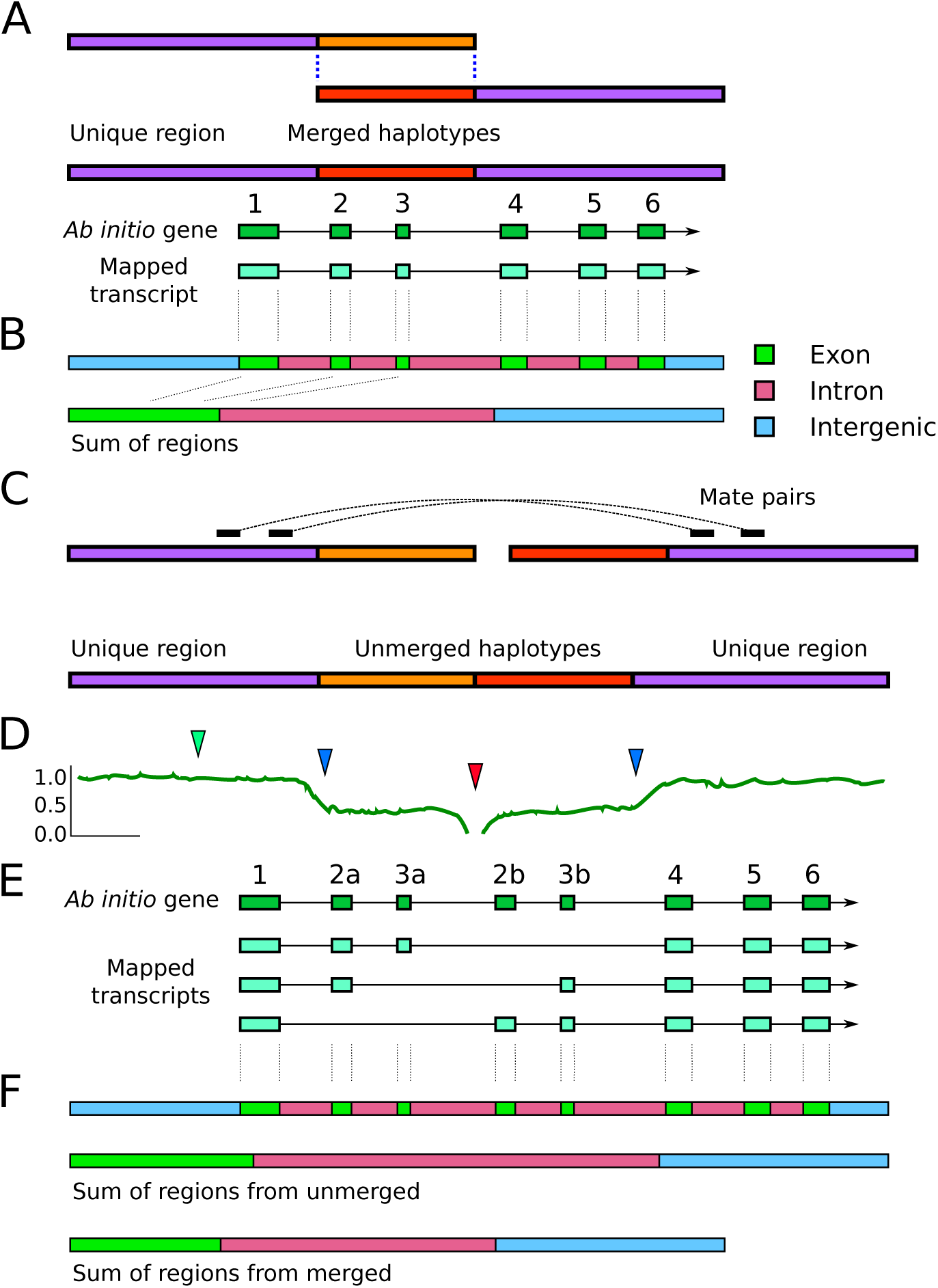
Schematic of misassembly and the effects on genic fraction analyses. (A) During assembly, regions that are heterozygous (differing by SNPs or indels) are combined to make a single reference contig. When genes are predicting that this locus, or when assembled transcripts are aligned to the genome, the correct exon structure is found. (B) Regions are defined as exon, intron, or intergenic, as in Fig 1. (C) Reference genomes are a mix of the maternal and paternal haplotypes, but not uniformly. Rather than being merged into a single sequence, highly heterozygous regions may be assembled as different contigs that get erroneously fused during scaffolding steps. Mate pairs that bridge the two purple unique regions will instead result in a head-to-tail joining of the two unmerged haplotype sequences. (D) Hypothetical plot of read coverage across the contig. The green arrow shows a region of normal coverage (1x) while the blue arrows show sites where coverage is reduced because reads for each haplotype map separately. At the fusion point between the two haplotypes (red arrow), no reads will map since the sequence is an artifact, or is represented by a gap. (E) Mapped transcripts (or ESTs) or transcripts derived from mapped RNAseq reads (such as by Cufflinks or StringTie) may only be mapped to one of the two haplotypes, thereby producing a staggered exon structure. A mapped transcript can only align to either exon 2a or 2b, but not both, likewise for 3a or 3b, yet all other exons are unique and would align correctly. Genes predicted *ab initio* may annotate both sets of exons (2a/3a and 2b/3b), which may result in a duplication in some part of the protein, or a premature stop codon if 3a and 2b are out of phase. (F) For this hypothetical case, the sum of 10 the regions would appear to have increased total exon size and the total intron size compared to the same genomic locus where the haplotypes were correctly merged.

Regions with relatively high heterozygosity may fail to be merged in this way, leaving contigs of both haplotypes in the assembly (Fig 3C). During subsequent scaffolding steps, contigs of separate haplotypes can be fused head-to-tail if mate pairs are bridging the unique regions. Because this head-to-tail joining is an artifact, no reads should map at the junction point, resulting in a region of zero coverage at the junction and flanked by regions where coverage is half of the expected value (Fig 3D). One additional feature may reveal this artifact: exons in the unmerged sections may be individually annotated but mapped ESTs or *de novo* assembled transcripts may show a staggered exon pattern (Fig 3E) because transcripts can only map to one of the two possible exons (2a or 2b, 3a or 3b). This may increase the ratio of intron:intergenic sequence (Fig 3F), but also falsely indicate that splice variation is more prevalent for this gene.

### Reannotation and changes following RNAseq reannotation

Keeping in mind the above error sources, some of the genomes used in our study had obvious problems of too much or too little genic content that would confound our analyses. For instance, the total amount of exons in the JGI annotation of *T. adherens* (Triad1) was only 14Mb, over twofold lower than the related species, the placozoan *H. hongkongensis*, and thus it was expected to contain many more or longer genes than were present in the original Triad1 annotation. Because of this, we remade a gene annotation for five of the species (see Methods) and used two additional publicly available annotations for *N. vectensis* and *A. queenslandica*. For most species, the reannotation dramatically increased the total amount of exons as well as the total bases of genes (Fig 4). The only exception was *B. floridae*, where the original annotation had predicted 90% of the genome as genes, while the reannotation had annotated only 44.8% as genes.

**Figure 4:**
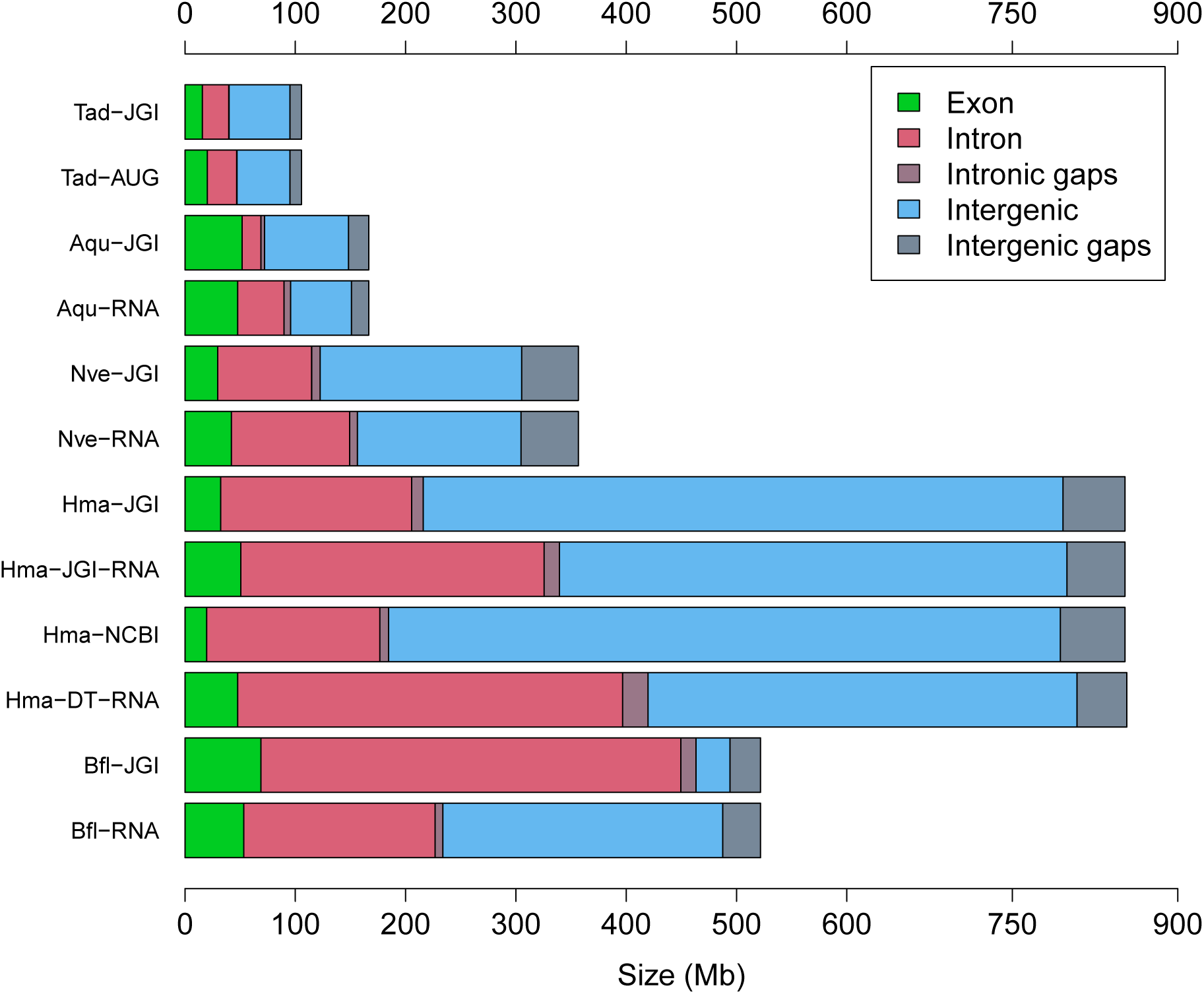
Proportions of exons, introns, and intergenic sequences. Barplot showing the summed proportions of genomes composed of exons (green), introns (red) and intergenic sequences (blue). The reannotation for *O. bimaculoides* was not shown for clarity, as this genome is substantially larger than the others. Abbreviations are as follows: Tad:*T. adherens*, Aqu:*A. queenslandica*, Nve:*N. vectensis*, Hma:*H. magnipapillata*, Bfl:*B. floridae*. JGI refers to the original annotations for each species downloaded from the JGI Genome Portal. RNA refers to reannotation (see Methods) with RNAseq. Hma-NCBI is the NCBI GNOMON annotation of *H. magnipapillata*. Hma-DT-RNA is the Dovetail reassembly of *H. magnipapillata* annotated with RNAseq. AUG is the reannotation using AUGUSTUS for *T. adherens*.

We then compared the ratio of intron:intergenic sequence across seven of the reannotated species (Fig 5). Across these species, reannotation significantly shifted the ratio of intron:intergenic sequence, approaching a 1:1 ratio (difference from 1:1 ratio, paired two-end t-test, p-value: 0.014). For *M. brevicollis*, the genome is very small and the majority is exons, so the reannotation was likely to change gene boundaries (separating run-on genes) rather than defining many new genes; our reannotation contains 10,864 genes compared to the 9,196 genes in Monbr1 “best models”.

**Figure 5:**
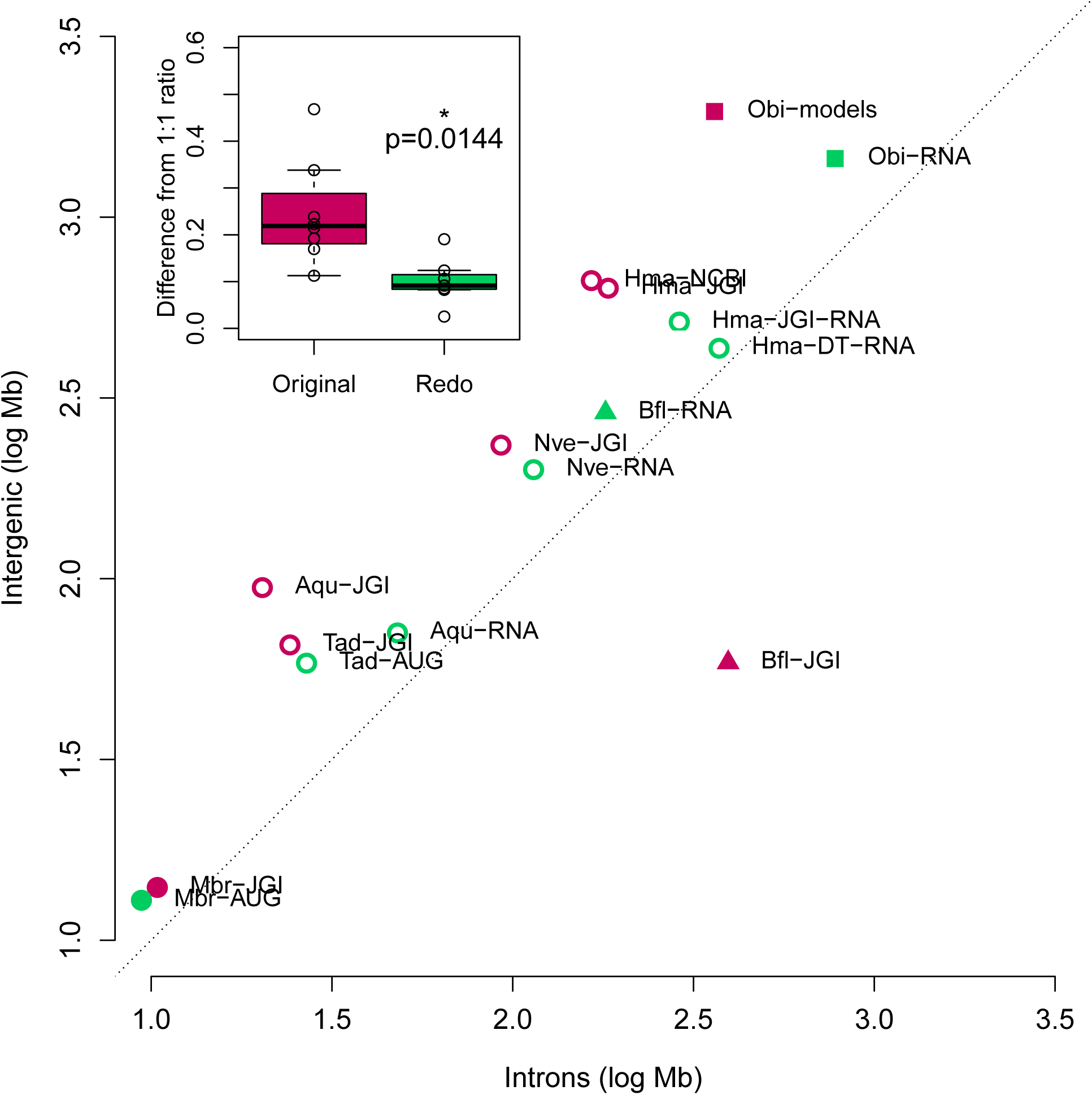
Improvements from reannotation. Log-scale plot of total intronic size versus total intergenic size where original annotations from the published genomes are shown in red and reannotations are shown in green. The dotted line shows a ratio of 1:1 as a reference. Abbreviations are as in Fig 4, with the addition of Mbr:*M. brevicollis* from the original JGI annotation and the redo with AUGUSTUS, and Obi:*O. bimaculoides* from the published gene models and the reannotation with Tophat/StringTie. The inset graph shows box plot of difference of the intron:intergenic ratio to 1, showing the reannotated genomes (green) are significantly closer than the original version (paired two-end t-test, p-value: 0.0144).

### Basic trends related to genome size

We observed linear correlations of total genome size to both total intronic size and intergenic size (Fig 6) (p-value: *<* 10^−37^ for both parameters). A much weaker correlation is observed for exons (R-squared:0.3856, p-value: 10^−8^). Because the total amount of exons in the largest genomes can be several times greater than the total size of the smallest genomes used in the study, a correlation is likely to be observed. Thus, the total amount of exons is necessarily affected by total genome size, even if this is not strongly correlated.

**Figure 6:**
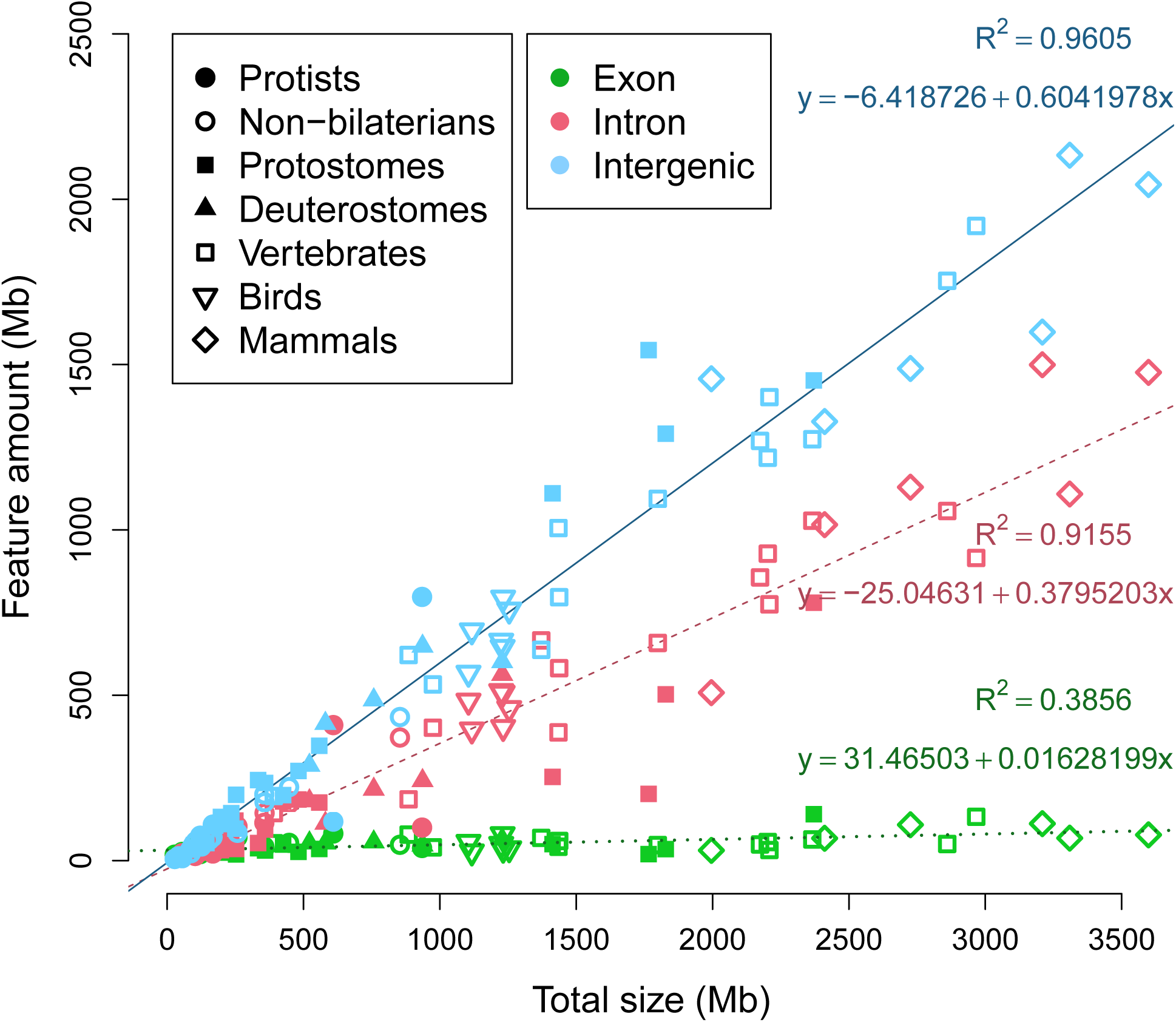
Comparison of features to total genome size. The sums of exons, introns, and intergenic regions are plotted against total genome size. Linear coefficients of determination of the three features are displayed by their respective lines. For legend symbols, Deuterostomes refers to all invertebrate deuterostomes, Vertebrates excludes Birds and Mammals.

### Average intron and exon length

The average length of introns linearly scales with the total genome size (Fig 7), in agreement with another study [18]. However, the average exon length is clearly constrained across animals relative to total genome size, and this may be related to interactions with nucleosomes [85]. Most species have an average exon length between 200 and 300 bases (mean of 263bp), higher than values reported from previous surveys of exon length [21, 86]. It must be stated that the average values presented here should not be taken as final, because variations in format of the annotations and quality of the genomes will affect the values. Since many genomes are only annotated with *ab initio* gene predictions, UTR exons may be missing from the annotation and all downstream calculations. Given that the first exon and intron tend to be longer than other exons and introns [21], respectively, absence of five-prime UTRs may result in an underestimation of the average exon length for that species.

**Figure 7:**
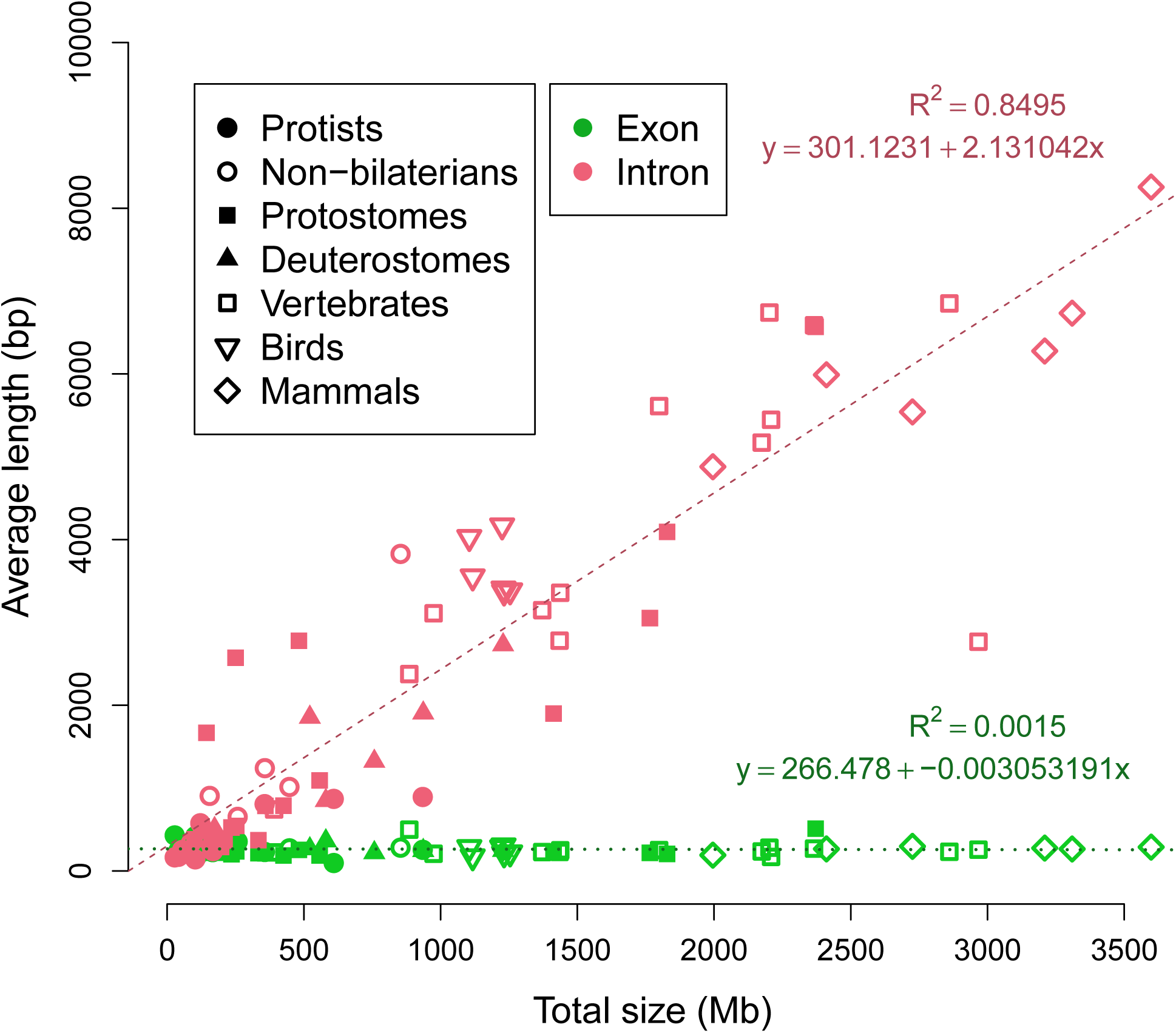
Average length of exons and introns. Plot of the average length of exons (green) and introns (pink) as a function of total genome size across all species in this study. Linear coefficients of determination are displayed next to the green (dotted) and red (dashed) linear fit lines, for exons and introns, respectively.

### Nature of the exonic fraction

Unlike introns or intergenic sequence, the total amount of exons does not show a strong linear correlation with total genome size (as seen in Fig 6). However, there is a hyperbolic correlation of the relative fraction of exons (megabases of exons divided by total megabases) compared to total genome size (Fig 8). The smallest genomes are dominated by exons, while the largest genomes are dominated by introns and intergenic regions. This implies a relatively fixed pool of exons or coding space that becomes spread over the genome as the total size increases. The hyperbolic trend resembled the observed hyperbolic relationship between total genome size and coding proportion [18]. As coding exons are a subset of total exons, measurements of total exons may be a reasonable approximation of coding sequence, but not necessarily vice versa.

**Figure 8:**
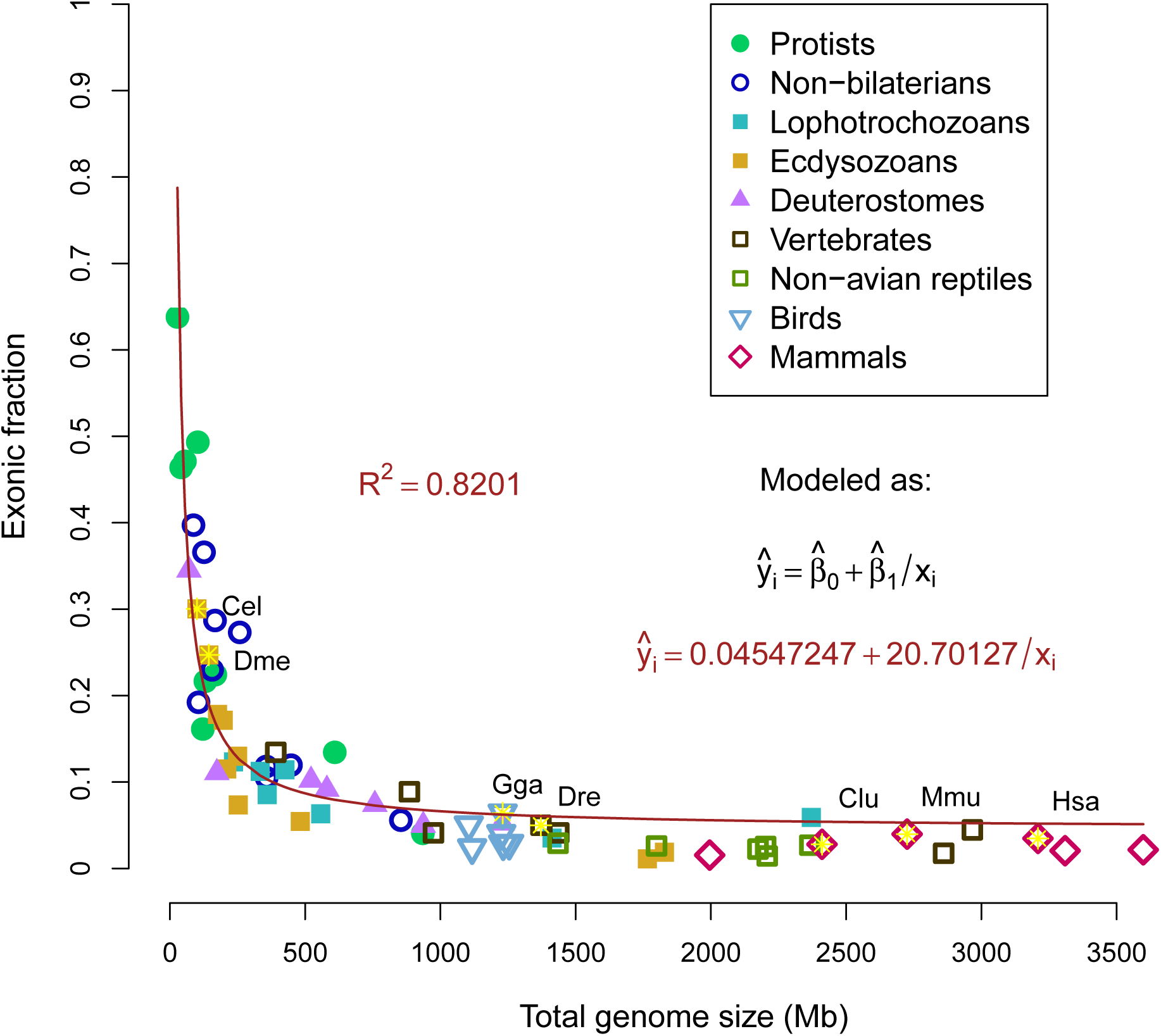
Exonic fraction compared to total genome size. Relative fraction of the genome that is defined as exons compared as a function of total size. Coefficients of determination of a hyperbolic model is displayed. Seven model organisms (human, mouse, dog, chicken, zebrafish, fruit fly and nematode) are indicated by three-letter abbreviations. The formula for the fitted model is displayed in red.

### Ratio of introns to intergenic

Because both intronic and intergenic fractions displayed a linear correlation to total genome size (Fig 6), we next examined the connection between the two fractions. While many species have a ratio of introns:intergenic approaching 1:1 (R-squared: 0.8286, p-value: 5.6 *\**10^−27^), the majority of genomes are composed of sequence annotated as intergenic regions (Fig 9).

**Figure 9:**
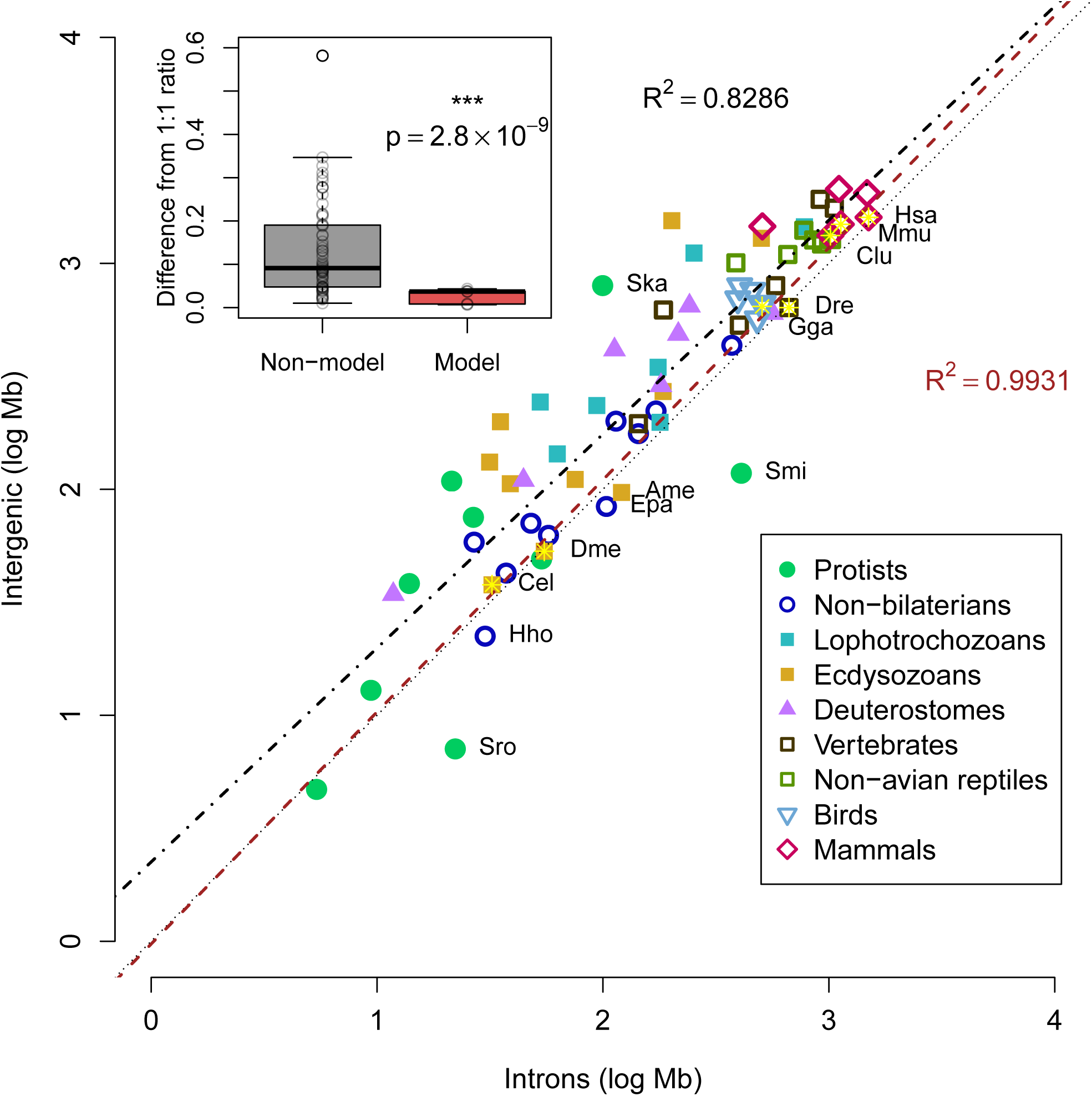
Comparing intronic and intergenic fractions. Log-scale plot of total intronic size versus total intergenic size. The dotted line shows a ratio of 1:1 as a reference, although most genomes are above this line. Seven model organisms (as in Fig 8) are indicated by three-letter codes with yellow stars. Black dashed line displays the linear fit of all species in the study (R-squared: 0.8286, p-value: 5.6 10^−27^), while the red line displays the linear fit for only the seven model organisms (R-squared: 0.9931, p-value: 1.3 10^−6^). Names are displayed for model species, two dinoflagellates (Ska:*S. kawagutii*, Smi:*S. minutum*) and select species with ratios of intron:intergenic greater than 1, choanoflagellate *S. rosetta* (Sro), honeybee *A. mellifera* (Ame), anemone *E. pallida* (Epa), and placozoan *H. hongkongensis* (Hho). All other species names are omitted for clarity. The inset graph shows box plot of difference of the intron:intergenic ratio to 1, showing the model organisms (red) have significantly different ratios compared to the rest of the genomes (paired two-end t-test, p-value: 2.8 *\** 10^−9^).

Because of the potential issue of gene annotation accuracy, we tested the linear correlation of introns:intergenic sequence for seven model organisms likely to have accurate annotations. A better linear fit was observed when restricted to the model organisms (R-squared: 0.9931, p-value=1.3*\** 10^−6^), suggesting that deviations from the 1:1 ratio of intron:intergenic sequence are due to missing annotations, rather than biological differences. Genomes of model organisms are significantly closer to the reference line (two-tailed t-test, p-value: *<* 10^−7^ for either absolute distance from 1:1 reference or absolute difference of intron:intergenic ratio to 1), suggesting that the better annotations of model organisms predict a ratio of 1:1 of intron:intergenic sequence. Overall, the comparison of genomes of model to non-model organisms is compatible with the hypothesis that the predicted amount of the genome that is transcribed varies more by annotation quality than biological differences.

We then examined if there is a difference between genomes of vertebrates and invertebrates. No significance difference is observed between the two model invertebrates and five vertebrates (two-tailed t-test, p-value:0.99). Among all species in the study, significant differences are tenuous and highly dependent on the species selected (Figure 10). For example, chordates against non-chordates is not significant (p-value:0.128) while vertebrates against invertebrates is significant (p-value:0.008). However, the observed significance appears to be an artifact of the abundance of low-quality genomes of protostomes, since comparison of vertebrates against non-bilaterians is not significant (p-value:0.83). This difference is most simply explained by the similarity between vertebrate groups. That is to say, annotation of a new mammalian genome is facilitated by existing knowledge of gene structures in other mammals, rather than true differences in genome organization.

**Figure 10:**
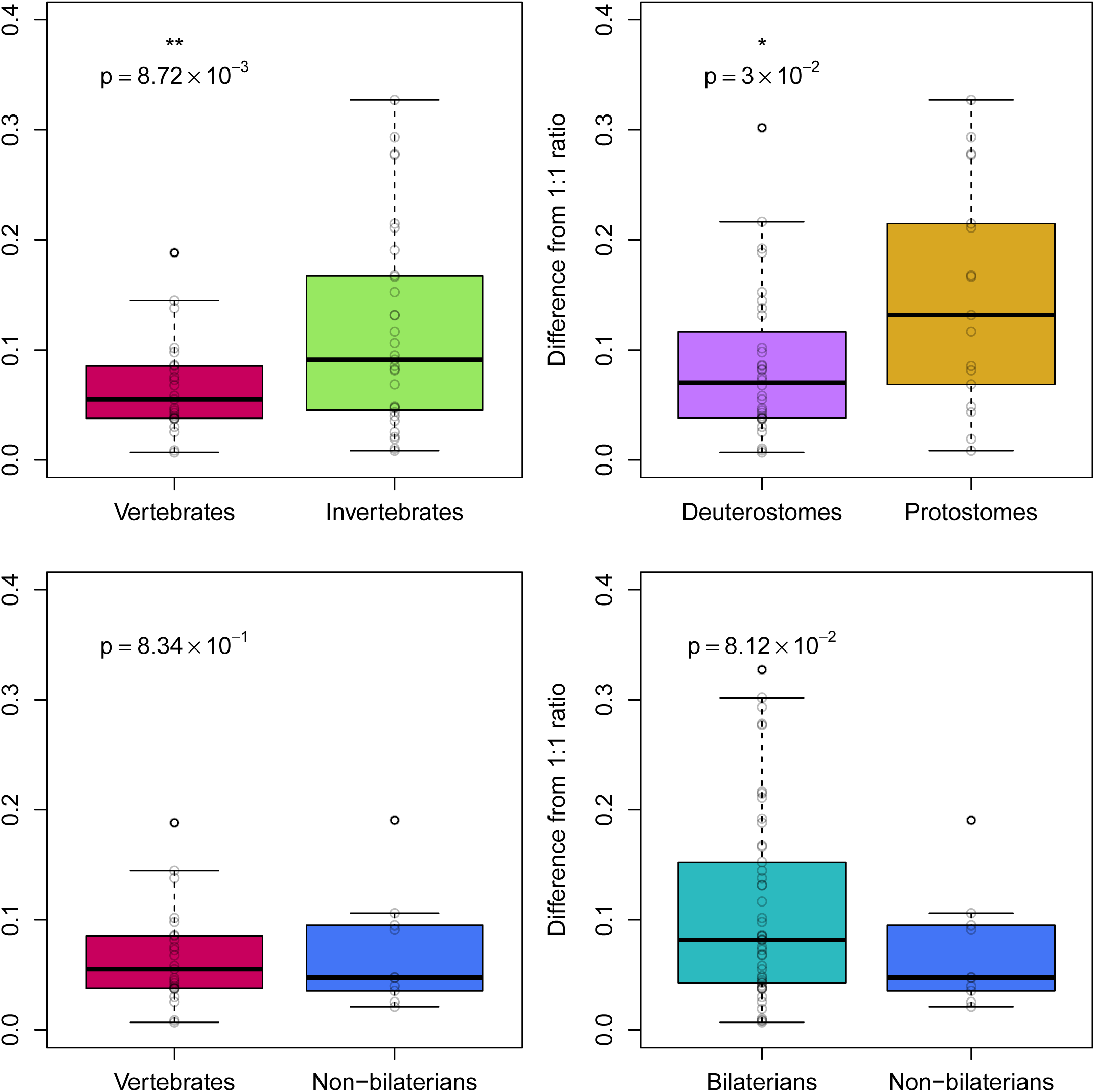
Comparing intron-intergenic ratios among animal groups. Difference of the intron:intergenic ratio to 1 across four pairs of animal groups. Invertebrates (green) includes all non-bilaterian taxa. Deuterostomes and protostomes are both assumed to be monophyletic.

Several genomes are below the 1:1 reference line, indicating slightly more introns than intergenic, such as the choanoflagellate *S. rosetta*, the honeybee *A. mellifera*, the anemone *E. pallida*, and placozoan *Hoilungia hongkongensis*. For *A. mellifera*, it was noted that improvements in versions of the genome also included better placement of repetitive intergenic sequences [71], suggesting that the relative surplus of introns is merely due to the absence of some intergenic sequences in the final assembly. As for *E. pallida* and *H. hongkongensis*, these species stand out as having relatively high heterozygosity, 0.4% [87] and 1.8% (manuscript in preparation), respectively. Although these values are lower than the observed heterozygosity in many other invertebrates [88], some highly heterozygous sequences may have caused assembly problems during scaffolding (as proposed in Fig 3).

### Evolution of the genic fraction

The amount of the genome that is composed of genes was highly variable across the genomes in our study, ranging from 12.5% up to 87.1% of the genome. Unlike the exonic fraction, the relationship of the fraction of the genome that is genes to the total size is less obvious (Fig 11), in part because this parameter is most subject to gene annotation accuracy. The fraction of the genome that is exons (and perhaps coding) appeared relatively fixed (Fig 8), yet the intron size was linearly correlated to the total size (Fig 6), therefore the fraction that is genes (exons and introns combined) was expected to be a combination of the two trends. Three correlation models were tested: hyperbolic (double-log), exponential (single-log), and linear. Of these, the hyperbolic model fit best (R-square: 0.3649, p-value: *<* 10^−8^), and no correlation was found for the other models. Restricting the linear model to only genomes larger than 500Mb found essentially no correlation (R-squared: 2.5 *\**10^−4^), suggesting that the genic fraction is unrelated to total genome size in large genomes but not in small genomes.

**Figure 11:**
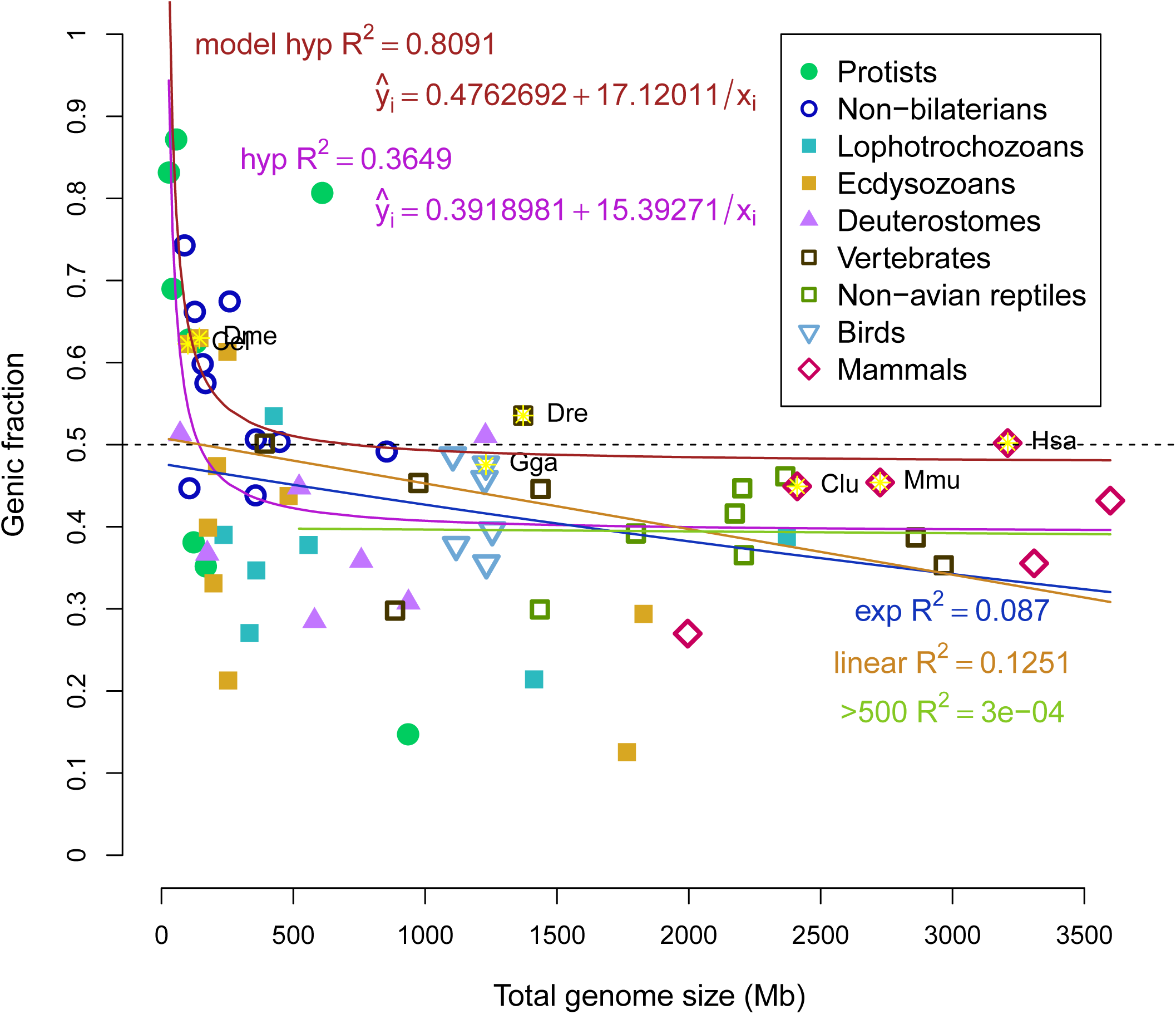
Genic fraction compared to total genome size. Relative fraction of the genome that is defined as genes compared as a function of total size. A number of correlative models (hyperbolic in purple, exponential in blue, linear in orange) were tested and coefficients are displayed. Linear correlation is expected to be zero if genic and intergenic fractions “expand” indifferently after a certain size, which appears to be around 500Mb. Linear correlation including only genomes larger than 500Mb is also displayed as the green line. Seven model organisms (as in Fig 8) are indicated by three-letter codes and yellow stars. The hyperbolic correlation model for the seven model organisms is shown in red. The formulae for the fitted models are displayed in red and purple, for model organisms and all organisms, respectively.

Again, the importance of gene annotation accuracy cannot be ignored and needs to be emphasized. When restricting to the seven model organisms, the range of values is narrower, from 44.9% to 62.9%. The same three correlation models were applied to the genomes of model organisms, again finding that the hyperbolic model best explained the variation in the genic fraction of model organisms (hyperbolic R-squared: 0.8091, p-value=0.0058; exponential R-squared=0.6709; linear R-squared=0.6835). Rather than simply having no correlation to total size, these results suggest that the genic fraction is fixed at around 50% in large genomes.

## Discussion

### Diagnostic relationship of introns to intergenic sequence

An increasing number of genomes of any non-model organisms are sequenced to answer evolutionary questions. For example, genomes of taxa from all four non-bilaterian groups were recently sequenced to understand how similar these genomes are to humans [8,24,33,34], and found that we share much more in terms of genes with these groups than had been previously thought. Yet, one of the main challenges in studying the genomes of non-model organisms is that there is little *a priori* information about gene structure or content. It would be expected that finding orthologs of human genes is relatively easy, but does not inform us about other genes that differ from humans. How should we know when we have found all of the genes? Our results provide some guidance here and suggest that there is a constant ratio of introns to intergenic sequence in all animals. This relationship holds even for animals with small genomes, such as the model organisms *D. melanogaster* and *C. elegans*, suggesting that organisms with small genomes and many currently sequenced invertebrates are subject to the same forces as organisms with large genomes.

### Unusual cases of genomes

Based on our model, the majority of genomes appear to be underannotated, in that substantial portions of the genome are not predicted to be transcribed when in fact many probably are. However, only two species, the lancelet *B. floridae* and the dinoflagellate *S. minutum*, display a dramatic trend in the opposite way, that is, the majority of the genome is annotated as genic (being primarily introns).

For the lancelet *B. floridae*, the original JGI gene models had annotated almost 90% of the genome as genes [23], the majority (85%) of that sequence being introns. Our reannotation of this genome displays the opposite trend, where more of the genome is intergenic than intronic. The original JGI annotations did not include any validation of the predicted genes, as predictions were made using mapped ESTs only as inputs for the gene model training. From this, we consider it more likely that the RNAseq-based transcripts more accurately resemble the true gene structures, albeit missing some genes. In addition, other evidence suggests that the *B. floridae* annotations may have been unusual or erroneous [89]. A study of domain combinations found that *B. floridae* had by far more fusions than any other species (across all eukaryotes) and had to be excluded from the analysis [90], precisely the expected result if the majority of genes were erroneously fused.

The only other species have a much larger ratio of intron to intergenic was the dinoflagellate *S. minutum*. It was described that its genome contained many long stretches of genes on the same strand, sometimes continuing for hundreds of kilobases [41]. The authors also note that the *de novo* assembled transcriptome appears to contain transcripts spanning multiple genes and containing multiple open reading frames, indicating the possibility that dinoflagellate symbionts can make cistronic transcripts. This species is not an animal, so it should not be assumed that animal modes of transcription are conserved across all eukaryotes. However, it should be noted that a recently published genome of another symbiotic dinoflagellate species *S. kawagutii* [40] does not display the same pattern, and instead appears to have a much greater fraction of intergenic regions than introns.

### Genome composition across metazoa

Previous studies have discussed problems with trying to relate the number of genes to the size of the genome [91–93]. One study [18] found a weak positive correlation between genome size and number of genes. This parallels our finding that total exonic sequence is weakly correlated to total genome size (Fig 6). However, this measurement can be problematic if the genome assembly is highly fragmented, containing a large number of short contigs or scaffolds. In such cases, gene number is unlikely to correlate to genome size for the same reason as the difficulties in predicting the genic fraction, that is, it is strongly affected by gene annotation errors. In our schematic (Fig 2), a gene that is split up onto three contigs would therefore be counted as three genes, albeit short ones. If this occurs on a genome-wide scale, the count of genes will be inaccurate. Parts of genes would be individually annotated as genes, increasing the total gene number without much change to the total number of exonic bases.

Rather than relying on counts of genes or determining coding sequence, we instead examined sequence that is annotated as exons. We found that while a weak positive correlation is observed between total exonic bases and genome size, most of the difference in size is related to introns and intergenic sequence. The amount of the genome that is composed of introns is linearly related to the total genome size (Fig 6). Also considering the measured linear correlation of intergenic sequence to total size, it is not surprising that most species have roughly a 1:1 ratio of introns:intergenic sequence (Fig 9). This appears to be the case regardless of genome size or the total exonic sequence. For instance, the genome of the choanoflagellate *M. brevicollis* has 9.3Mb of introns and 10.1Mb of intergenic sequence (a ratio of 0.92) compared to 19.3Mb of exons.

Therefore, model animals (and probably all animals) transcribe nearly half of the genome, where species with smaller genomes (exon-rich) transcribe more than half (Figure 11). There does not appear to be a significant difference in the genic fraction based on animal group (Figure 10), that is, all animals appear to follow this rule. One study had shown that some larger metazoan genomes were depleted in genes [94], yet this study made use of a small number of species for comparison and included several chordates known for their very small genomes, the tunicate *C. intestinalis* and the pufferfish *T. rubripes*. The authors examined windows of 50kb and found that 80% of the human genome was lacking any gene [94], though it is unclear if this analysis was restricted to protein coding genes. However, we found that 50.2% of the human genome is composed of genes (93% of that is introns).

While genomes of the model organisms and many non-models organisms appear to follow the hyperbolic relationship of genic fraction to size, nonetheless, a large number of the genomes in this study appear to be composed of much less than 50% genes. That observation is best explained by the hypothesis that many genomes are missing genes. These missing genes may or may not be coding, though perhaps missing gene content is made of lineage-specific proteins. Because annotation of the genome by RNAseq per se cannot distinguish coding genes from non-coding ones, we could not determine coding fractions for all species. Even for putative non-coding transcripts, some may be coding [95–97], thus protein sequencing may reveal the true nature of these transcripts.

### Evolution of genomes

The genic fraction has a hyperbolic relationship to the total genome size. The modeled curve flattens around 500Mb, after that point, introns and intergenic regions are expected to expand, on average, equally across the genome resulting in approximately 50% of the genome as genes (the majority of that being introns) and the other 50% as intergenic sequence. It should be noted that larger genomes still have more exonic bases than small genomes, though the difference in total genome size across animals is mostly from introns or intergenic sequence.

It has been theorized that changes in genome size are a balance between short deletions and long insertions [98]. If the last common ancestor of all metazoans had a relatively small genome (under 100Mb, resembling some single-cell eukaryotes in our study), then the majority of modern animals have undergone dramatic expansion of their genomes, meaning dominated by insertions or duplications. How does this expansion occur and does it favor a novel origin of introns or expansion of intergenic sequences? Following the trend in Fig 9 and Fig 11, it appears that small genomes are dominated by genes because they are mostly exons, and both genes and intergenic sequences are expanded in equally as the genomes enlarge. Mechanistically, these insertions are likely to be mediated by transposable elements or replication errors. As small genomes become invaded by transposable elements (perhaps following some genomic stress like genome duplication), introns appear and expand at roughly the same rate as intergenic sequences producing a 1:1 ratio of intron:intergenic across all species (Fig 9).

Above a certain size (around 500Mb), genic and intergenic sequences expand almost equally, where 50% of the genome is genic; exons comprise an almost negligible fraction of the genome, which is otherwise composed of approximately equal fractions of introns and intergenic sequences. This might be explained by changes in diversity of transposable elements, as the highest diversity was found in genomes ranging from 500Mb to 1.5Gb [17]. Larger genomes appeared to be flooded by transposable elements of a single type. Thus, above 500Mb, it can be predicted that select transposable elements become prevalent and multiply throughout the genome, but on average end up expanding introns and intergenic sequences equally.

### Relationship to phenotypic complexity

The size of the genome can vary greatly even for closely related organisms. This has been called the “c-value paradox” [1, 99], based on the observation that although the many organisms have larger genomes relative to similar species (bigger “c-value”), this measurement does not equate with more or less complex organisms in a straightforward way. A classic example of this is frog genus *Xenopus*, where the genome of the species *X. laevis* is almost twice as large as the species *X. tropicalis* [100], though the animal is not twice as “complex”. Similar observations have been made that the number of genes appears unrelated to the size of the genome and the complexity (sometimes called the “g-value paradox” [91, 101]).

If neither genome size nor gene number are clearly related to complexity, then what is? Another relationship has been proposed between the usage of alternative splice variants and organismic complexity because variation in splicing can increase the number of potential proteins from an overall fixed pool of exons [102]. Vertebrates and specifically mammals tend to splice transcripts more than invertebrates (meaning models fruit fly and nematode) [103, 104]. One study reported a good correlation (R-squared of 0.80) of splicing to organismic complexity measured by cell types [105], but also reported that this trend effectively disappeared when correcting for sequencing depth, using the number of ESTs available as a proxy for annotation quality. The largest invertebrate genome used in that study was the deer tick *I. scapularis*, which did have a measured number of cell types but unfortunately could not be analyzed further, leaving the bulk of the analysis weighted heavily by mammals and small-genome insects.

However, other studies report that alternative splicing is more frequent when the surrounding introns are long [106, 107], suggesting that organisms with large genomes (and therefore larger introns) might be predisposed to splice. This could suggest that some of the invertebrates in our study may have more complex splicing patterns than are annotated in the current genome versions. For the largest invertebrate genome in our study, the octopus *O. bimaculoides*, only 14.8% of loci appeared to have alternative splice variants [45]. In our reannotation we found only 6.4% of all loci have any type of splice variant. However, the majority of predicted transcripts (75%) are single exon loci, and possibly many genes are fragmented across multiple contigs (as in Figure 2). When restricted to loci with multiple exons (15% of total loci), 41% have more than one variant. These data from *O. bimaculoides* suggested that overall patterns in splicing do not display a reliable connection to organismic complexity when complexity is generalized across animal groups. However, without proper measurements of cell types from the octopus, it cannot be assumed that the number of cell types resembles the value for the fruit fly, which was implicit in other studies given that all protostomes were effectively represented by insects [105]. Thus, it could be the case that the octopus, with a large genome, has a large number of cell types and many genes are spliced, all in agreement with the splicing-complexity hypothesis.

It is a challenge to separate these observations from biases in sequencing depth (of transcripts or ESTs) and data availability. In our study, we could only make use of five invertebrates with relatively large genomes, the cnidarian *H. magnipapillata*, the pearl oyster *P. fucata*, the horseshoe crab *L. polyphemus*, the deer tick *I. scapularis*, and the octopus *O. bimaculoides*. On the other hand, NCBI has over 100 genomes of mammals available for download. Alternatively, the repertoire of splice factors or the genes that are most spliced may be of greater importance than just splicing in general. Our understanding is likely to be improved with more deeply-sequenced transcriptomes from large-genome invertebrates.

### Limitations

Because we were making use of mostly public data, our analyses were subject to both technical and biological limitations. There are a small number of taxa with sequenced genomes from many invertebrate groups. Because the majority of sequenced vertebrate genomes are large and the majority of sequenced invertebrate genomes are small [92], the axis of simple invertebrate to complex vertebrate is synonymous with small to large genomes, and thus the prevalence of splicing in large-genome animals may be a consequence of the size of the genome and complexity may be only correlated. This issue is not simple to resolve, as there may not be members in all animal groups with both small and large genomes. For instance, a survey of genome sizes across Porifera stated that the largest genome out of the 70 species sampled was around 600Mb [108]. Thus, there may not be any “large” genomes in this phylum, and likewise for other invertebrate groups. Compared to birds, however, where the smallest genome identified to date is from the black-chinned hummingbird (estimated 910Mb) [109], perhaps no bird will be found that has a “small” genome.

Our use of public genome annotations was limited in part from difficulties in defining elements. Much like definitions of transcribed pseudogenes, the identification of long-intergenic non-coding RNAs, or lincRNAs, presents a paradox of definitions. Non-coding RNAs with known functions are arguably genes, such as the X-inactivation transcript Xist, thus any functional transcribed intergenic RNA is by definition not intergenic; it is genic. This distinction rests upon discovery of a function of these putative RNAs. In the context of the ENCODE project or MouseENCODE [110], transcription was found of intergenic regions accounting for almost another 20% of the genomes of human and mouse, depending on the analysis [111, 112]. If this were all functional, then the genic fraction of the genome would be far above 50% for large genomes and the ratio of intron:intergenic sequence would not be expected to be close to 1:1. Alternatively, if most of these intergenic transcripts are non-functional “noise”, then our results are supported as presented. Therefore, consideration of the importance or genic quality rests upon the distinction between functional RNAs and noisy transcription. Existing data are not adequate to identify functions, but several experiments may improve our understanding. Conceptually, the most straightforward approach is knocking out regions of transcribed “gene deserts” in mouse or human cells, but on a larger scale than a previous study [113]. Additionally, better models of transcriptional noise or random transcription may inform whether or not the observed transcriptional patterns from the ENCODE project are consistent with noise.

## Conclusion

We have shown that a set of animals from 12 phyla transcribe at least half of their genomes in a size-dependent fashion. For large genomes, the amount of exons is almost negligible, where introns account for most of the genic sequence. In such cases, genic sequence is almost equal to the amount of intergenic sequence. Whereas for small genomes, exons can be a major fraction of the genome, resulting in the appearance of gene-dense genomes. This parity between introns and intergenic sequence is likely a universal feature of animal genomes, though this may be tested with addition of many more animal taxa from other phyla that do not have sequenced members. Previous findings of genomic differences between animal groups are likely to result from a sampling bias, rather than biological differences. Future improvements in assembly and annotation of animal genomes may reveal unanticipated sources of complexity and gene regulation with implications for the evolution of animals.

## Acknowledgments

W.R.F would like to thank M. Eitel for helpful comments on the manuscript. This work was supported by a LMUexcellent grant (Project MODELSPONGE) to G.W. as part of the German Excellence Initiative. The authors declare no competing interests.

